# The demographic history of African *Drosophila melanogaster*

**DOI:** 10.1101/340406

**Authors:** Adamandia Kapopoulou, Susanne P. Pfeifer, Jeffrey D. Jensen, Stefan Laurent

**Author notes:** Author for Correspondence: Stefan Laurent, MPIPZ, Cologne, Germany.

## Abstract

As one of the most commonly utilized organisms in the study of local adaptation, an accurate characterization of the demographic history of *Drosophila melanogaster* remains as an important research question. This owes both to the inherent interest in characterizing the population history of this model organism, as well as to the well-established importance of an accurate null demographic model for increasing power and decreasing false positive rates in genomic scans for positive selection. While considerable attention has been afforded to this issue in non-African populations, less is known about the demographic history of African populations, including from the ancestral range of the species. While qualitative predictions and hypotheses have previously been forwarded, we here present a quantitative model fitting of the population history characterizing both the ancestral Zambian population range as well as the subsequently colonized west African populations, which themselves served as the source of multiple non-African colonization events. These parameter estimates thus represent an important null model for future investigations in to African and non-African *D. melanogaster* populations alike.

## Introduction

Populations of *Drosophila melanogaster* span five continents, making this organism a widely utilized system to study patterns of local adaptation. Yet, this complex underlying demographic history represents unique challenges for disentangling non-neutral from non-equilibrium processes (*e.g.* Jensen et al. 2005; Teshima et al. 2006; Thornton & Jensen 2007; Pavlidis et al. 2010), and thus numerous studies have worked to better illuminate the correct demographic null model. Considerable effort has been made in understanding the species’ expansion to Europe (*e.g.* Thornton & Andolfatto 2006; Li & Stephan 2006), Asia (*e.g.* Laurent et al. 2011), and the Americas (*e.g.* Kao Joyce Y. et al. 2015).

However, it is only in the past decade that African demographic history has been similarly scrutinized. In one of the earliest studies, Dieringer Daniel et al. (2004) surveyed X-chromosomal microsatellite variation from thirteen sampling locations across Africa, describing considerable population structure between North, West, and East Africa. Pool & Aquadro (2006) surveyed nucleotide variation at four 1-kb fragments in 240 individuals from sub-Saharan Africa, and described a distinct East-West geographic pattern, suggesting that western Africa may have been recently colonized from the East. Simultaneously, Li & Stephan (2006) examined dozens of non-coding X-chromosome regions from a population sampled in Zimbabwe, suggesting strong evidence of population growth. In a much larger-scale study, Pool et al. (2012) sequenced whole-genomes from 139 wild-derived strains from 22 sampling locations in sub-Saharan Africa. Based on levels of variation and *F*_ST_, they qualitatively described a fit to a model in which Zambia represents the species origin, with subsequent population expansion, structuring and gene flow across the continent – though they concluded on the need for proper demographic model fitting in order to better elucidate these patterns. In addition, Singh et al. (2013) examined a 2Mb region in 20 individuals sampled from Uganda, also finding support for population expansion, but also suggested an associated population bottleneck out of the initial ancestral range (presumably being Zambia, hundreds of miles to the south).

Following this important work, we here focus our study on Zambia as the likely population of origin, and West Africa as a likely source of multiple widely studied non-African populations (Figure 1). We quantify the demographic history of these regions, including the timing of West African colonization, effective population sizes, and rates of gene flow (Supplementary Figure 1). Furthermore, given known segregating inversions as well as the associated difficulties that may arise if they are left unaccounted for, we have carefully curated a dataset for the purposes of inferring these underlying neutral demographic parameters, which may serve as the basis for future studies.

**Figure 1:**
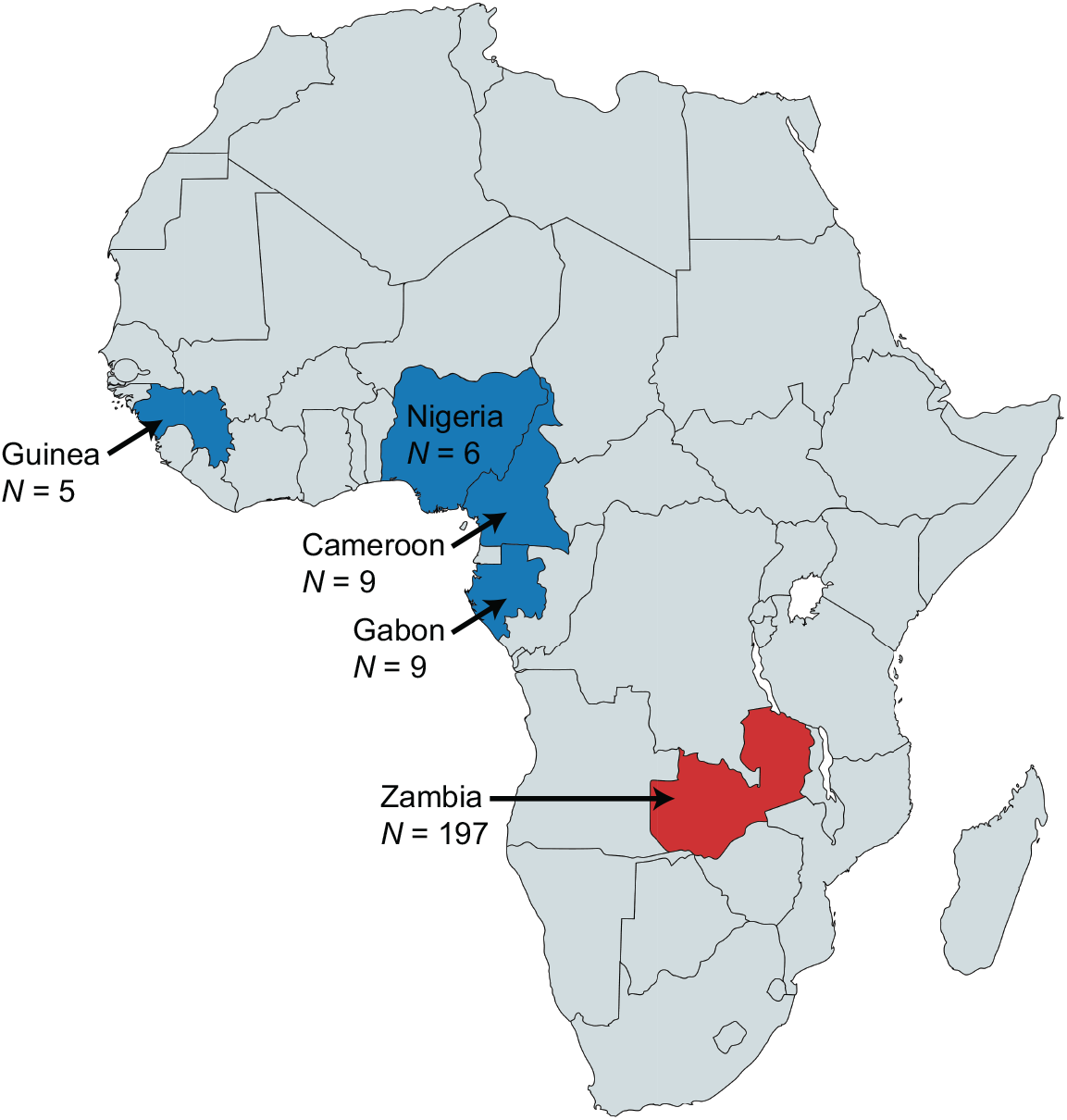
Geographic distribution of the five *D. melanogaster* populations. Samples (sample sizes indicated by *N*) were obtained from the Phase 2 (blue) and Phase 3 (red) of the Drosophila Population Genomics Project (Pool et al. 2012; Lack et al. 2015).

### Inferring Population History

The levels of genetic differentiation between individuals were assessed using a principal component analysis. The first principal component, explaining 2.7% of the variation, separates the Zambian individuals from the West African individuals, which cluster according to their sampling location (*i.e*., Cameroon, Gabon, Guinea, and Nigeria; Supplementary Figure 2). In contrast, Zambian individuals cluster in two distinct groups based on chromosomal inversions carried by the individuals (Supplementary Figure 3). This pattern was well described by Corbett-Detig & Hartl (2012) who noted that polymorphic inversions in *D. melanogaster* affect genomic variation chromosome-wide, with trans-effects beyond the inversions’ breakpoints. To avoid the confounding effects of these segregating inversions on subsequent demographic inference, 121 Zambian individuals carrying at least one inversion (*i.e*., In2RNS, In2Lt, In3R, and In3LOk) were excluded from any further analyses.

Population structure was then assessed using an admixture model to infer individual ancestry proportions using *sNMF* (Frichot & François 2015), a statistical method to evaluate the ideal number of ancestral populations. The best-fit model (*i.e*., the model with the lowest minimal cross-entropy) had two ancestry components (Figure 2a), strongly supporting the division of individuals from Zambian and West African populations, with evidence of admixture between them (Figure 2b). Principal component analysis confirms the two population clusters inferred by *sNMF*, with no additional sub-genetic stratification of the Zambian individuals (Figure 2c).

**Figure 2:**
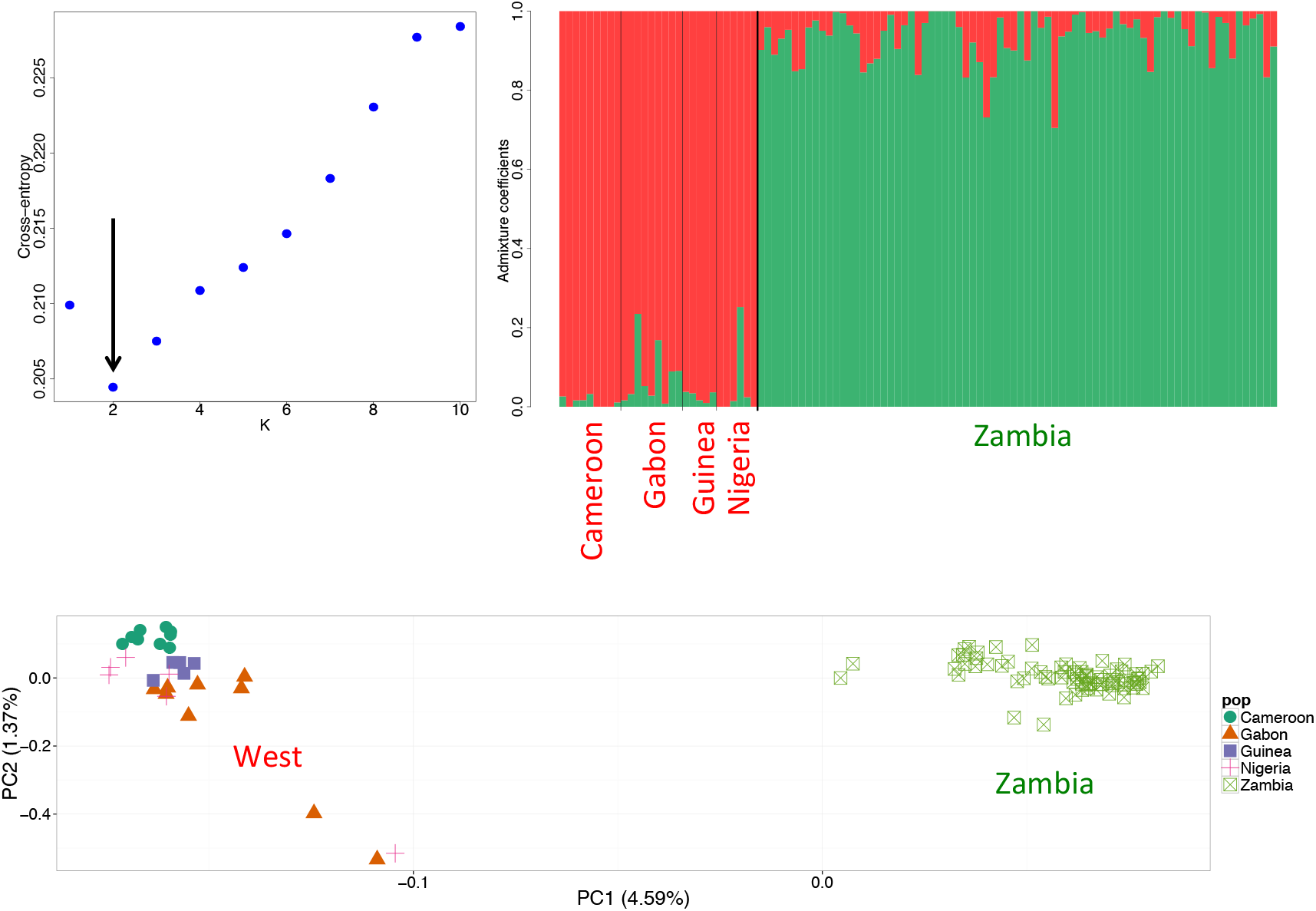
Genetic structure of African *D. melanogaster* populations. a) The number of *K* ancestry components best explaining the data was assessed by calculating the cross-entropy corresponding to the model. The best-fit model (*i.e*., the model with the lowest minimal crossentropy) had two ancestry components (*K*=2). b) Individual admixture proportions. c) Principal component analysis (symbols correspond to individuals from different populations; green square: Zambia (*N*=76 individuals which do not carry the chromosome arm’s specific inversion); green circle: Cameroon (*N*=9); orange triangle: Gabon (*N*=9); purple square: Guinea (*N*=5); red cross: Nigeria (*N*=6)). Data was thinned to prune for linkage, excluding SNPs with an r2>0.2 within a 50 SNP window. Percentages indicate the variance explain by each principle component.

Given the observed population structure, the demographic history of Zambian and West African populations was investigated using six different two-population demographic models, allowing for both size changes as well as gene flow among the populations. Three of the six models assumed that populations remained at a constant size with either no gene flow, symmetric migration, or asymmetric migration between them (Supplementary Figure 1). To account for the fact that West African populations exhibit lower nucleotide diversity levels than populations from south-central Africa (*π*= 0.0086 in Zambia, *π*= 0.0077 in West Africa; and see Pool et al. 2012; Lack et al. 2015), suggesting a potential population bottleneck during their recent colonization from the ancestral range (Haddrill et al. 2005), the remaining three models allowed for population size changes (Supplementary Figure 1). The demographic model best fitting the data (Figure 3; Supplementary Table 1) inferred a reduction in population size followed by exponential growth for both the Zambian and West African populations after their split around 70kya, with on-going gene flow. In addition, the parameter estimates obtained for the ancestral and present effective population sizes (*N_e_*(anc) = 1,525,061 (95% CI: 1,498,713 − 1,562,754); *N_e_*(Zambia) = 3,160,475 (95% CI: 2,933,313 − 3,447,248)) reiterate the higher levels of variation observed in the putative ancestral range of the species.

**Figure 3:**
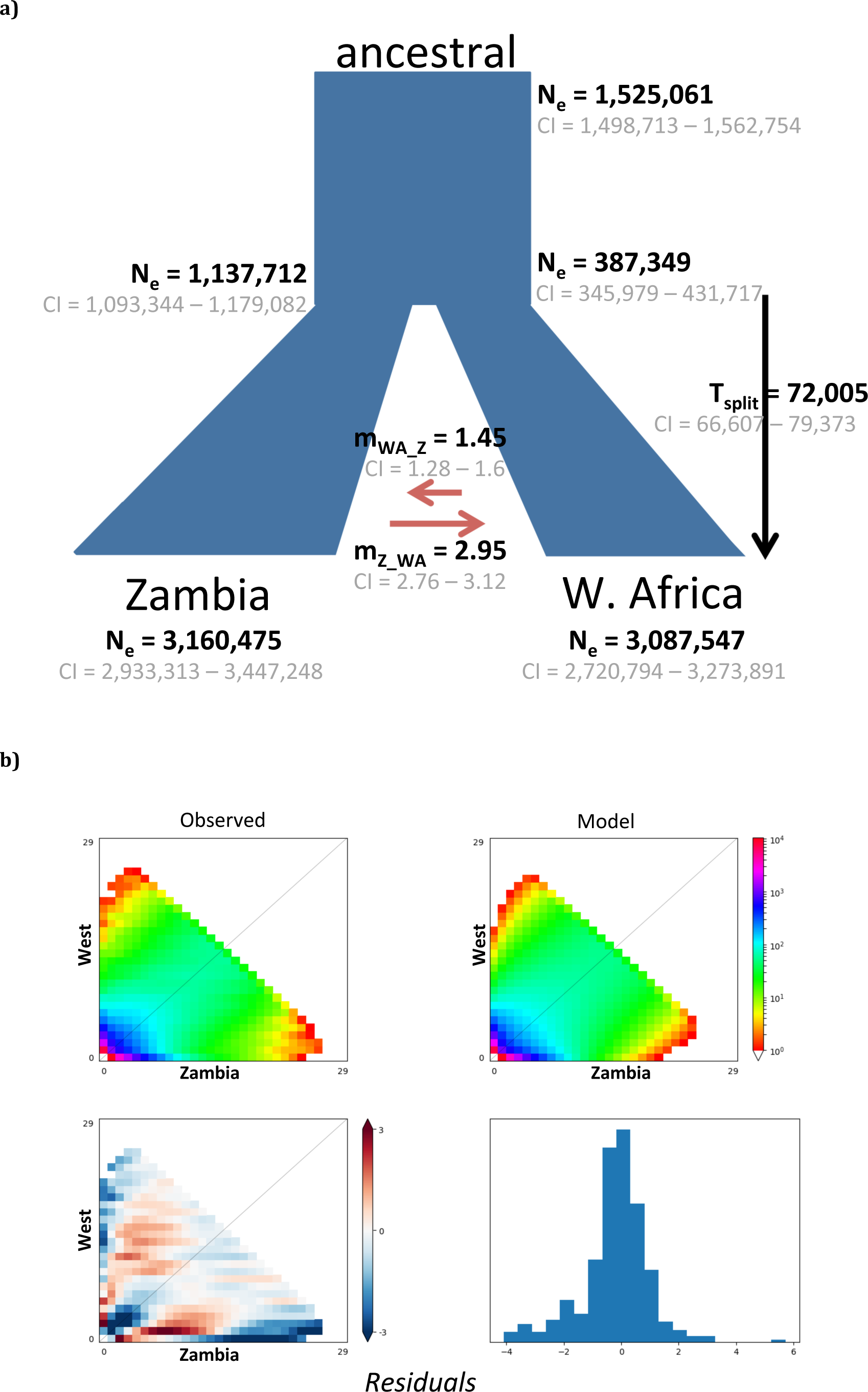
Parameter estimates inferred by ∂a∂ i under the best demographic model. a) At time *T_split_*, the ancestral population splits into two distinct populations, which grow exponentially with asymmetric migration (*m*) between them. The time of the split (*T_split_*) was estimated in generation times, which were converted to years, assuming ten generations per year (Laurent et al. 2011). Effective population sizes (*N_e_*) for the ancestral, West African, and Zambian populations were directly estimated by fixing the mutation rate (*μ*) to 1.3 × 10^−9^ per base pair per generation (Laurent et al. 2011). 95% confidence intervals (CI) were calculated for each parameter estimate by generating 150 parametric bootstrap replicates of the best model. Note that the mode of the bootstrapped parameter estimates corresponds approximately to the obtained maximum likelihood value estimate. b) Comparison of Joint SFS for the observed data (left) and the best model (right). Below are shown the residuals of the model.

While the specific parameter values inferred are of particular importance for explicitly modelling an appropriate demographic null in future studies, and represent the first estimates of split times between the ancestral range and West Africa, the qualitative patterns are largely consistent with previous supposition. Namely, the population structure (Pool & Aquadro 2006), and effective population sizes (Laurent et al. 2011), as well as the underlying growth and colonization models themselves (Pool et al. 2012), are all largely in agreement with previous studies.

### Concluding Thoughts

In concordance with Corbett-Detig & Hartl (2012), we have demonstrated the ability of inversions to create significant sub-structure within a single population sampled from a single location, potentially confounding downstream demographic inference. Indeed, we find that even when polymorphisms within the inversion breakpoints were not considered in the analysis, the signature persists and is visible when analyzing other markers located on the same chromosomal arm (Supplementary Figure 3). By removing these individuals from the analysis, and by carefully curating the dataset for neutral sites, we have quantified the demographic histories characterizing these sampling locations. We find evidence for strong growth in populations inhabiting both regions, consistent structure separating West Africa from Zambia, as well as evidence for on-going gene flow particularly in the direction of south/central to west. Thus, this well-fit non-equilibrium demographic model of both the ancestral range of the species as well as the source population of subsequent non-African colonization events, represents a uniquely appropriate null model for future investigations pertaining to the demographic and adaptive histories of both African and non-African populations of *D. melanogaster*.

## MATERIAL AND METHODS

### Samples

Publicly available whole-genome sequence data from haploid *D. melanogaster* embryos originating from Guinea (*N*=5), Nigeria (*N*=6), Cameroon (*N*=9), Gabon (*N*=9), as well as from Zambia (*N*=197) was obtained from the Phase 2 and Phase 3 of the Drosophila Population Genomics Project (DPGP) (Pool et al. 2012; Lack et al. 2015, 2016), respectively (Figure 1). Specifically, genomes previously aligned to a common *D. melanogaster* reference sequence were downloaded from the Drosophila Genome Nexus (DGN) (Lack et al. 2015, 2016) and variants on both arms of chromosome 2 (*i.e*., chr2L and chr2R) and chromosome 3 (*i.e*., chr3L and chr3R) were identified using the SNP-sites C program (Page et al. 2016).

As chromosomal inversions may be targeted by natural selection in *D. melanogaster* (Corbett-Detig & Hartl 2012), known inversions were excluded from all demographic analyses (information on inversion breakpoints was obtained from the DGN (Lack et al. 2015; http://www.johnpool.net/Updated_Inversions.xls). To further minimize the confounding effects of linked selection on demographic inference, the dataset was limited to putatively neutral regions of the genome, including four-fold synonymous degenerate sites (Grenier et al. 2015) as well as the 8^th^ to the 30^th^ base of introns smaller than 65bp (Parsch et al. 2010). The resulting dataset contained 82149 variants.

### Inferring Population Structure

Population structure was investigated using two methods, which cluster individuals based on their genetic similarity using a set of independent SNPs (*i.e*., SNPs with an *r*^2^>0.2 within a 50 SNP window were excluded from the dataset using PLINK v1.07 (Purcell et al. 2007)). Evidence of population structure was assessed using both a principal component analysis (PCA) as well as the *sNMF* function implemented in the R package LEA v2.0.0 (Frichot & François 2015). The latter implements an admixture model (Pritchard et al. 2000; Patterson et al. 2006) which uses sparse non-negative matrix factorization to infer individual ancestry proportions based on *K* potential components. Using a cross-validation technique, *K* values ranging from 1 to 10 were examined, and, following (Frichot et al. 2014), the best *K* was selected to minimize the cross entropy.

### Demographic Inference

The demographic history of south-western African *D. melanogaster* populations was inferred from the distribution of minor allele frequencies (*i.e*., the folded joint site frequency spectrum) obtained from the putatively neutral segregating sites using ∂a∂i 1.7.0 (Gutenkunst et al. 2009), a diffusion approximation method. Given the genetic differentiation between populations, six different two-population scenarios (corresponding to samples originating from West Africa - *i.e*., Guinea, Nigeria, Cameroon, and Gabon, as well as Zambia) were tested, allowing for both population size changes as well as gene flow among the populations (Supplementary Figure 1). Thereby, gene flow was modelled either as symmetric or asymmetric, and considered only between the time of the population split and the present.

For every demographic model, 10 independent runs were performed using different starting points and the parameter estimates for the best run (*i.e*., the estimation with the highest likelihood) reported. 95% confidence intervals (CI) were calculated for each parameter estimate by generating 150 parametric bootstrap replicates of the best model. Effective population sizes (*Ne)* were directly estimated by fixing the mutation rate (μ) to 1.3 × 10^−9^ per base pair per generation (Laurent et al. 2011). Generation times were converted to years, assuming ten generations per year (Laurent et al. 2011). The best-fitting demographic model was selected based on the Akaike’s information criterion (AIC) score (Akaike 1974).

## ACKNOWLEDGEMENTS

We thank Roman Arguello for helpful discussions and for providing the coordinates of short introns and four-fold degenerate coding sites for the neutral set of loci. We also thank Athanasios Kousathanas and Anna-Sapfo Malaspinas for sharing their population genetics and statistical knowledge with us. This work was supported by grants from the Swiss National Science Foundation and the European Research Council to JDJ.

